# Statistics of cortical representational drift can enable robust readout

**DOI:** 10.1101/2024.09.26.614914

**Authors:** Charles Micou, Timothy O’Leary

**Affiliations:** Department of Engineering, University of Cambridge, United Kingdom

## Abstract

Representational drift of fixed stimuli, learned tasks and familiar environments is observed in many brain areas, leading to reconfiguration of population codes over days to weeks. This raises the question of whether downstream brain regions employ mechanisms to track changes in population activity and thus preserve the fidelity of the information they extract. We show that the statistical properties of drift have a significant impact on such mechanisms. Over an extended period, a net change in population tuning due to drift can arise from an accumulation of small changes distributed across the population, or via abrupt jumps that affect smaller subsets of cells at each time point. We demonstrate that an adaptive readout can exploit the heavy-tailed statistics of abrupt jumps to maintain a more stable readout using a simple inference mechanism. Using experimental data, we investigate the extent to which heavy-tailed drift statistics are observed during representational drift in the posterior parietal cortex and visual cortex. We find that experimentally measured drift does not conform to a Gaussian random walk. Instead, we find sudden jumps in neural tuning that would be advantageous for a downstream observer adapting to changes in representation. These observations motivate future study to determine whether adaptive decoding mechanisms exist in the brain and to determine the physiological mechanisms that shape the statistics of representational drift.

## Introduction

A principal function of the brain is to form a model of reality. The neural circuits that underpin this model do not have direct access external world. Instead, neural populations must build up and maintain representations of stimuli, environments and events through statistical regularities in upstream activity. However, these statistics are subject to change, which presents an inference problem: did a change occur due to an event in the external world, or due to some internal change in upstream neural circuitry?

Extensive work has shown that while some component of ongoing change can be regarded as mean-reverting, stationary fluctuations or noise, a significant component appears to be nonstationary over a timescale of days to weeks. Such nonstationary changes are expected if the conditions of a laboratory task change, or if an animal’s performance in a task changes. However, a wealth of studies find evidence of neural representations of familiar environments and mastered tasks changing without any obvious external evidence of learning or change in behaviour, a phenomenon known as representational drift [1]. Representational drift has been observed in a variety of brain regions: in the hippocampus [2, 3, 4], the visual cortex [5, 6], the posterior parietal cortex [7], the auditory cortex [8], and the olfactory cortex [9].

Representational drift generically leads to a gradual degradation in the accuracy of a fixed readout of information encoded in population activity, eventually rendering that readout no better than chance (Fig. 1.a) [10]. This motivates the idea that neural populations must adapt to changes in the neural circuits they interact with in order to continue exchanging a stream of useful information from an encoding that is being continually redistributed over individual neurons [11, 12, 13]. How the brain implements this adaptation remains an intriguing puzzle. Existing work has proposed separately that feedback mechanisms can externally drive correction to readouts [10], that simple Hebbian learning is sufficient to adapt to drift [14, 15] and can be supplemented by spontaneous reactivation to strengthen representations [16], and that homeostatic plasticity either in isolation [17] or combined with Hebbian learning [18] can produce stable readouts without the need for an explicit error-correction signal.

**Figure 1.**
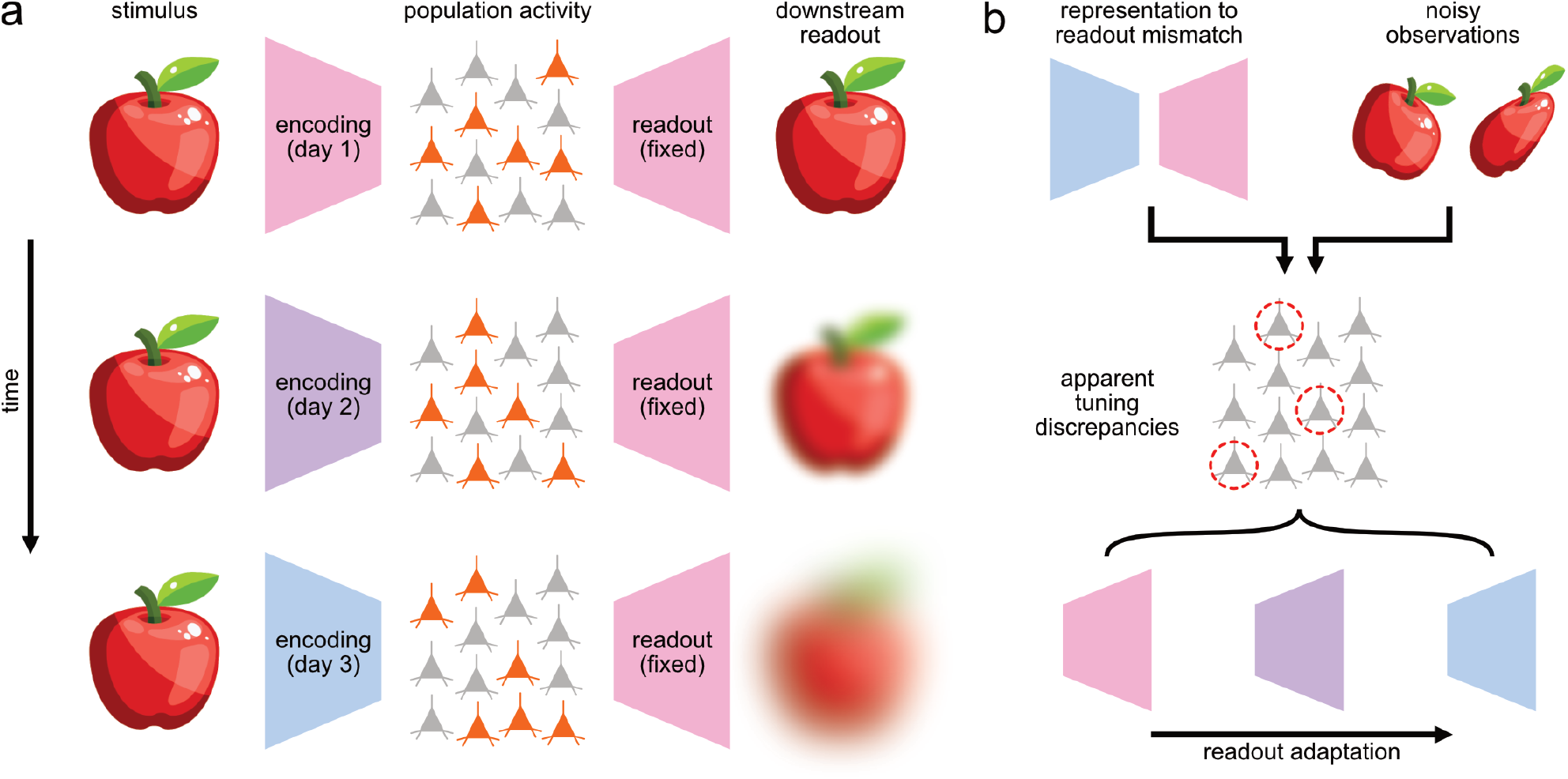
**(a)** A neural representation describes how patterns of population activity can encode a stimulus. A readout correctly matched to the encoding can infer the stimulus from population activity. However, as the representation drifts, the fixed readout becomes increasingly mismatched, and the quality of the downstream readout gradually deteriorates. **(b)** A downstream readout can attempt to correct for drift by identifying discrepancies in activity that suggest changes in tuning, but must take care not to confuse inherently noisy observations of the world for nonstationary changes in representation.

Our central argument in this work is that while such adaptation strategies are tractable (Fig. 1.b), qualitative statistical properties of the changes in tuning taking place at the level of individual neurons play a crucial role in determining both the maximum drift rates that can be compensated for and the amount of noise in observations the strategy can tolerate.

To support this argument, we first present a first-principles computational model of a method to stably extract an accurate signal from a drifting population long after a fixed readout would have degraded beyond the point of usefulness. We describe this method in terms of an unsupervised ‘adaptive decoder’: an algorithm that can infer changes in how stimuli are encoded in a population. We emphasise that our formulation is not intended to be a faithful description of an algorithm implemented by the physiology of the brain, but instead as a best-case normative example of such a mechanism.

We use this adaptive decoder, which is constrained to rely only on statistical regularities in activity to update its parameters, to illustrate fundamental limitations in the extraction of underlying representational changes from noisy observations, and how these limitations are mitigated by heavy-tailed statistics. We then demonstrate that the advantages of these heavy-tailed statistics present a design trade-off: compensating for drift without these heavy tails requires either larger, more redundant neural populations, or the rates of drift must be slower.

Finally, we explore whether biology makes use of these advantageous statistics by considering two existing and previously studied in vivo datasets that exhibit representational drift.

The first dataset, originally published in Driscoll *et al*. (2017) [7], contains recordings from the posterior parietal cortex (PPC) in mice during expert performance of a delayed choice task in a virtual T-maze [19]. This work originally measured drift by considering a sparse encoding of position along the main arm of the maze, with mean neuron firing forming peaks at specific locations, and measuring the changes in peak location. Recording sessions are spaced one day apart, and drift is visible between individual sessions.

The second dataset, originally published in Marks & Goard (2021) [6], contains recordings from the V1 region of the visual cortex in mice during exposure to both orientated grating stimuli and movies of naturalistic scenes. Instead of measuring drift through explicitly parameterised tuning curves, this work analysed clusters of neurons with comparable activity and used changed in membership of those clusters to estimate drift rates. This drift manifests more slowly than in the PPC dataset, and is measured in sessions spaced a week apart.

We characterise the distributions of tuning changes within both of these cortical populations and find evidence for heavy-tailed statistics, especially under high rates of drift.

## Results

### Adaptive decoding via continual tuning parameter inference

Drift is distinguished from simple noisy observation by its nonstationary statistics and resulting gradual degradation of fixed readouts. Drift is distinguished from the other form of nonstationary changes in neural populations, learning, by a constant representation fidelity: drifting representations do not represent the same variables more accurately (e.g. by increasing potential decoder accuracy), more efficiently (e.g. by using fewer neurons), or by using different overall encoding strategies (e.g. moving from a sparse to a dense encoding).

The objective of an adaptive decoder is to produce an accurate readout from the population in spite of drift. Each day, the adaptive decoder updates its estimates of how individual neurons are tuned to stimuli. Outside of initialisation, the adaptive decoder cannot ‘cheat’ with knowledge of stimulus values: it must instead use changes in the statistical regularities of the population to infer changes in the tuning of individual neurons.

Concretely, we model a neural population of *N* neurons by parameterising their responses **x** ∈ ℝ^*N*^ to a given stimulus **s** ∈ ℝ^*K*^ by defining *P* tuning curve parameters per neuron. The complete tuning of the population is therefore described by ***θ*** ∈ ℝ^*N*×*P*^. Individual neurons, subject to independent random noise *ξ*, have their responses defined by an activation function *a* common to all neurons in the population that relates tuning parameters to activity:

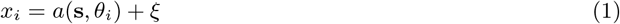

On the first day of an experiment, *t* = 1, we estimate of the tuning parameters 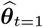 of the population by taking *M* samples of neural activity and the stimulus (**X**_*t*=1_ ∈ ℝ^*N*×*M*^ and **S**_*t*=1_ ∈ ℝ ^*K*×*M*^ respectively). On subsequent days *t* = 2, 3, …, *T*, the tuning parameters ***θ*** change due to drift and the ground-truth values of the stimulus are withheld from the adaptive decoder. The objective of the adaptive decoder is to maintain an accurate estimate of 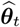 in spite of these restrictions (Fig. 2.a). For a set of tuning parameters ***θ***, we can use the activation function *a*, a prior distribution for stimulus exposure, and a model of the noise process for *ξ* to construct the function *g*, an approximation of the likelihood of *M* observations of neural activity **X**.

**Figure 2.**
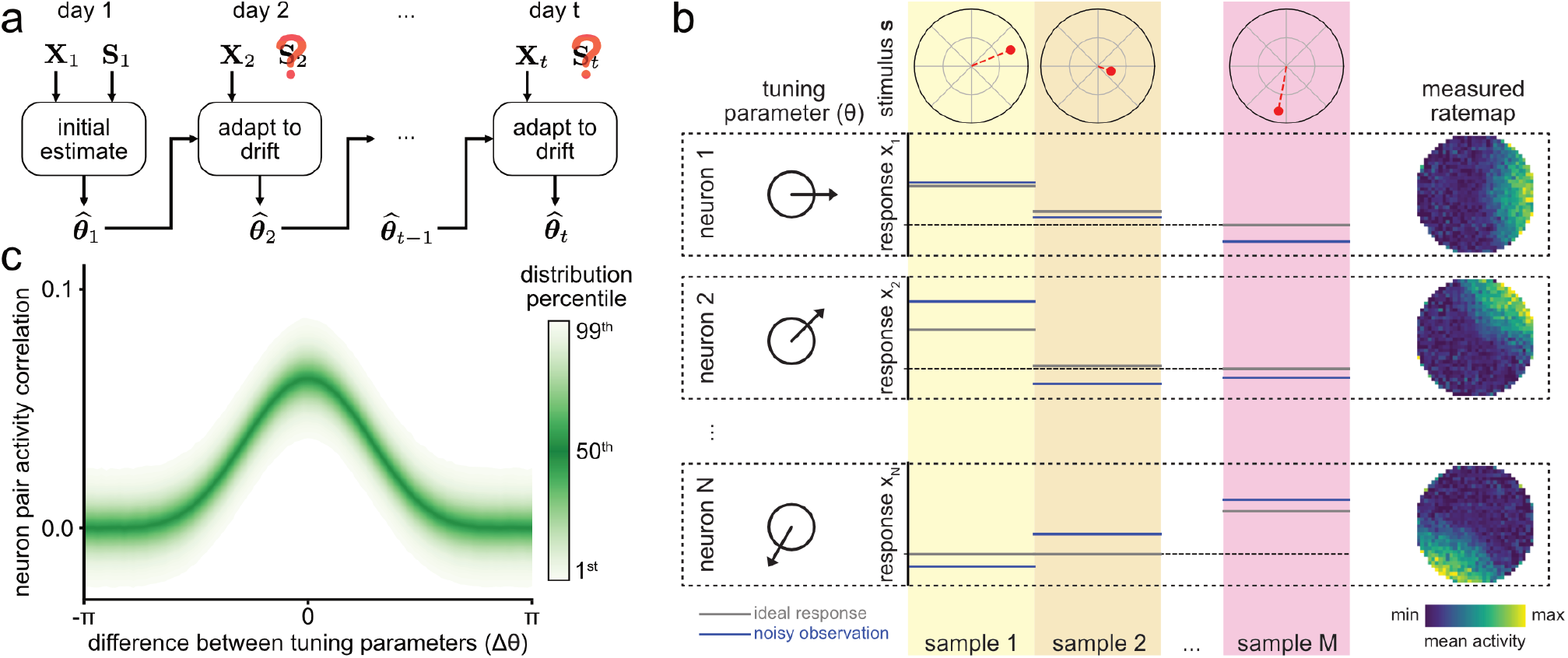
**(a)** Knowledge of the stimulus ground-truth from an initial experimental session is used to initialise an estimate of the tuning parameters that describe the relationship between stimulus and population activity. Subsequently, and with no further reference to the stimulus ground-truth, an adaptive decoding scheme attempts to keep these estimates accurate while the underlying tuning parameters drift. **(b)** A model of a neural population that encodes the orientation and magnitude of a stimulus. Each neuron in the population has a tuning parameter that describes its preferred orientation. The population is sampled multiple times during a session, with the activity of each neuron independently affected by noise at each sample. **(c)** Visualisation of the probability distribution for the value of the correlation between a pair of neurons as a function of the difference between their tuning parameters (simulation parameters *M* = 10^4^ and *σ* = 1.0).

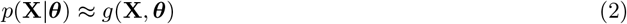

Naïvely, we might attempt to recover 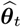 from **X**_*t*_ directly by finding the parameters that maximise *g*(**X, *θ***). However, because the activation function *a* is common to all neurons, the solution-space for maximising *g*(**X, *θ***) has symmetries that render the solution-space degenerate. Because *a* describes all neurons as interchangeable, a set of parameters maximising *g*(**X, *θ***) could be shuffled and assigned to entirely different neurons without affecting the solution: without their history, neurons lack identity.

By considering the tuning of neurons on the previous day, ***θ***_*t*−1_, we are able to break these symmetries. We use a stochastic model of how drift perturbs the population from one day to the next, *p*(***θ***_*t*_|***θ***_*t*−1_). This produces a maximum a posteriori update rule for tuning in the population.

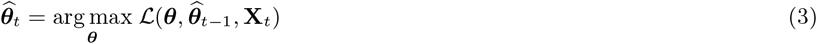

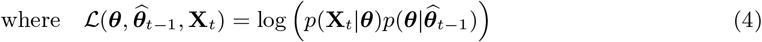

Treating drift as a stochastic process that is independent of neuron identity and in which each neuron drifts independently from other neurons, we can express *p*(***θ***_*t*_|***θ***_*t*−1_) in terms of a per-neuron drift prior *f*.

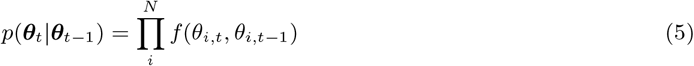

We then combine our model for observed activity *g* with our description for drift *f* and express the daily update rule for 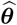 as:

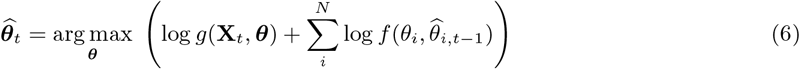

### Statistical regularities in the activity of a simulated population

As an illustrative example of this adaptive decoding technique, we consider a simple model of a neural population that encodes the direction and the magnitude of a stimulus: a system that is reminiscent of in vivo populations both of head-direction coding in the hippocampal formation and the response of populations in the visual cortex to grating orientations (Fig. 2.b). The population responds to a stimulus that corresponds to a point within the unit circle, defined by a magnitude and direction (i.e. *K* = 2), both of which are drawn from uniform distributions:

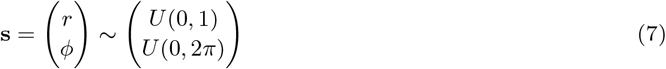

Neurons each have a single tuning parameter (*P* = 1), their preferred stimulus angle, and are subject to additive Gaussian noise.

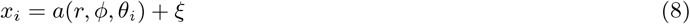

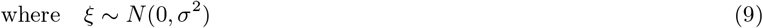

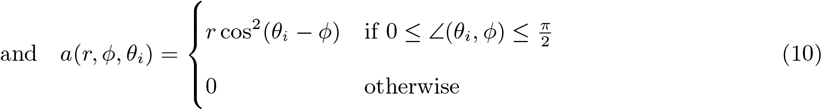

Where for the convenience of dealing with the circular geometry of this example, we have defined ∠(*ψ*_1_, *ψ*_2_) as the magnitude of the smallest angle between the angles *ψ*_1_ and *ψ*_2_. In this simple example, the mean decoding error in heading direction is directly equivalent to the mean error in the estimate of the tuning parameters.

To construct an expression for *g*(**X, *θ***), we consider pairs of neurons and the correlation of their activities within a session. Neurons with similar tuning values are more likely to be highly correlated with each other (Fig. 2.c). The correlation between neurons of a given angular separation is a probability distribution governed by two underlying random processes: the *M* random samples observed on any given day, and the observation noise *ξ* perturbing those observations (Fig. S1.a).

Taking *M* to be sufficiently large (Fig. S1.b) that the expected value of the correlation between two neurons of an angular separation Δ*θ* is well-approximated by:

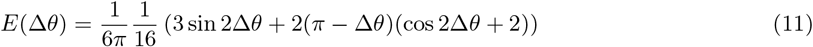

The distribution of the residual between the expected value and the measured value is normally distributed (Fig. S1.c). This allows a simple expression for the likelihood of the correlation a pair of neurons **x**_*i*_ and **x**_*j*_ with tuning parameters *θ*_*i*_ and *θ*_*j*_ observed *M* times within a session:

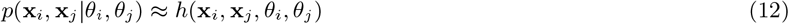

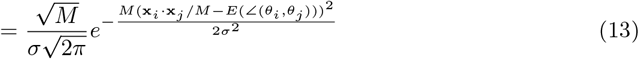

Treating the probabilities of pairwise correlations as independent, which is an approximation of *p*(**X**|***θ***), we can then construct an expression for *g*(**X, *θ***):

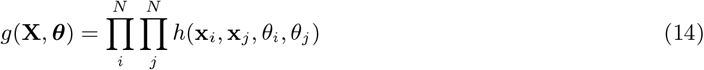

When solving the optimisation for 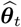, this amounts to minimising the sum of squared differences between observations of the correlations between neuron pairs and their expected values ***E*** as parameterised by ***θ***.

### Contrasting statistics of drift at the level of individual neurons

In our illustrative model the need to break symmetries using a drift prior is obvious: while *g*(**X, *θ***) helps determine the angular differences between neuron tuning parameters, it provides no information about the absolute orientation of any tuning parameter. Knowledge of how the population was tuned on an earlier day, as well as a drift prior describing how that tuning is likely to change, is required to anchor the system to an absolute frame of reference. Different models of drift can produce very similar degradation in the accuracy of a fixed decoder while having very different statistics at the level of individual neurons. We consider two simple but quite opposite models for drift in individual neuron tuning: a ‘gradual’ model, in which all neurons undergo relatively small tuning changes from one day to the next, and a ‘sudden’ (or ‘heavy-tailed’) model, in which most neurons retain their tuning while a small subset retune by making relatively large changes to their tuning (Fig. 3.a, Fig. 3.b).

**Figure 3.**
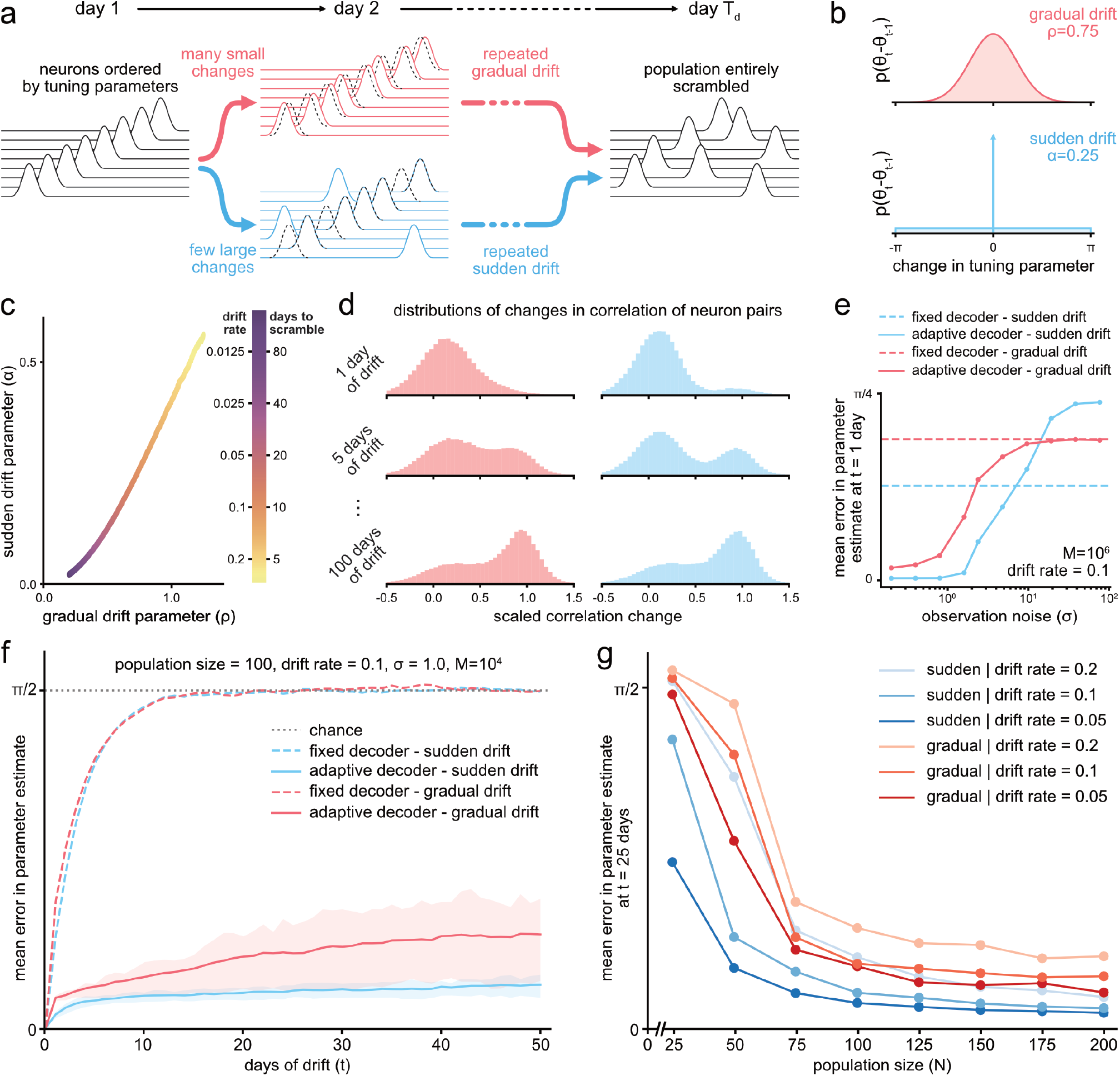
**(a)** Different underlying stochastic processes at the level of individual neurons, such as either gradually altering all neurons slightly or altering only a few neurons dramatically, can both produce drift. **(b)** These two example models of drift can be expressed as probability distributions describing the change in tuning parameter from one day to the next. **(c)** Equivalence between the two drift models determined by the expected number of days required to drive the error of a fixed decoder to within 5% of chance. **(d)** Histograms for the distribution of scaled changes in correlations between neuron pairs for both the sudden and gradual models of drift, shown for a relatively slow drift rate of 0.02. Only neuron pairs with a correlation of at least 0.05 on the latter day of comparison are displayed. **(e)** The resulting error in tuning parameter estimates after the first day of drift, shown as a function of the observation noise level *σ* for both gradual and sudden drift models. **(f)** Comparison of the accuracy of parameter estimates between fixed and adaptive decoders for both sudden and gradual drift models, both using drift rates of 0.1. Shaded regions show the interquartile range over 100 simulations. **(g)** The estimation error after 25 days shown for a variety of drift rates (all sufficiently high to scramble the population before 25 days) and for a variety of population sizes. Each datapoint represents the mean of 100 simulations.

In the gradual drift model, each parameter *θ*_*i*_ moves around the unit circle as if buffeted by Brownian motion: it is described by an Ornstein-Uhlenbeck process. The new value of a tuning parameter measured on a subsequent day is parameterised by *ρ*, the expected absolute distance drifted.

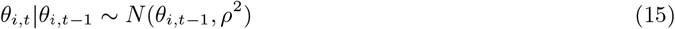

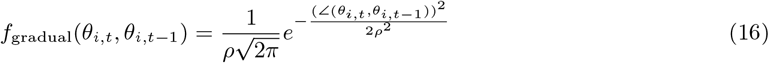

In the sudden drift model, each neuron undergoes a spontaneous retuning instead of gradually adjusting its tuning parameter. On any given day, it either retains its previous tuning parameter or selects an entirely new value. This distribution is parameterised by the spontaneous retuning probability *α*.

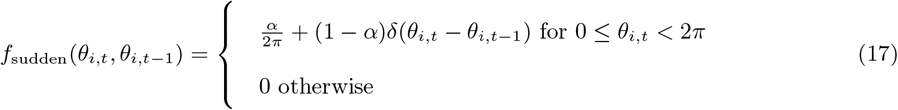

Where *δ* denotes the Dirac delta function.

To allow a fair comparison between sudden and gradual drift models, we define the drift rate of two models to be equivalent when the repeated application of drift scrambles the output of a fixed decoder in the same expected number of days *T*_*d*_ (Fig. 3.a).

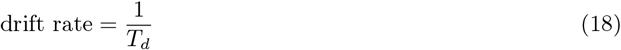

In the case of our illustrative example, we quantify the change in an individual neuron’s tuning between day

1 and day *t* as the angular distance between tuning parameters ∠(*θ*_1_, *θ*_*t*_). As drift progresses, the expected value of the change in tuning approaches the expected value between the initial tuning parameter *θ*_1_ and one chosen entirely at random *θ*_*r*_.

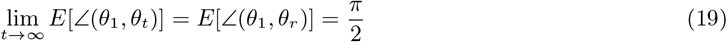

We describe a population as ‘scrambled on day *T*_*d*_’ if it satisfies:

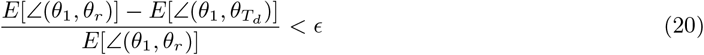

Which allows us to generate an equivalence between the parameters *ρ* and *α* of the two drift models (Fig. 3.c). We choose *ϵ*= 0.05, noting that the equivalence is not sensitive to this choice of parameter for small *ϵ* (Fig. S2.a, Fig. S2.b).

While the two models of drift provide identical long-term behaviour, the differences in short-term, transient behaviour have the potential for material impact on our adaptive decoding strategy. A small change in tuning might easily be confused for a noise-driven fluctuation (due either to the observation noise *σ* or a small number of samples *M*), while a large change in tuning is more distinctive. To visualise this effect, we consider the distribution of changes in the correlation between pairs of neurons. We define the scaled change in correlation across two days *D*_1_ and *D*_2_ as:

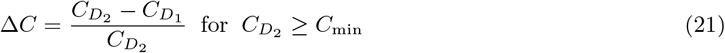

To aid the visualisation of the distribution we include only neurons with some small minimum correlation *C*_min_ on *D*_2_, as most neurons are not correlated with each other, which introduces a positive bias to the distribution. When we compare these distributions for the sudden and gradual models of drift (Fig. 3.d), we notice that over short time separations the sudden drift model is more distinctively bimodal. A single day apart, most measured changes in pairwise correlation are driven by statistical fluctuations: the component of the distribution near Δ*C* = 0.0. However, some neurons that were previously uncorrelated with each other have become correlated, accounting for the component of the distribution away from the origin. In contrast, under gradual drift it is more difficult to distinguish whether an individual change is due to a statistical fluctuation or a nonstationary change.

### Sudden drift is advantageous for adaptive decoding

In simulations of using the adaptive decoding scheme in our illustrative model, there is an obvious discrepancy between gradual and sudden drift in the difficulty of distinguishing statistical fluctuations from nonstationary changes. On the very first day of drift, the adaptive decoder can accurately infer changes in the tuning parameters of the population for both types of drift under low observation noise *ξ* (as parameterised by *σ*). However, as *σ* increases, adaptive decoders make larger errors when inferring tuning parameters subject to gradual drift (Fig. 3.e).

Adaptive decoders remain accurate long after fixed decoders perform no better than chance. However, because the strategy for estimating parameters relies on using estimates from earlier days, errors can compound and eventually degrade adaptive decoders (Fig. 3.f).

As the number of neurons in the population increases, adaptive decoders maintain their accuracy for longer for two reasons (Fig. 3.g). First, while additional neurons requires estimating an increasing number of parameters, the introduction of those neurons provide more neuron pairs with which to narrow the distribution of *p*(**X**|***θ***). The number of additional parameters to estimate grows with *N*, but the number of pairs available grows with *N* ^2^, which reduces the likelihood of poorly estimating the tuning of neurons. Second, additional neurons provide redundancy that is relevant over the course of many days of drift: it is statistically inevitable that, eventually, the tuning parameter of an individual neuron will be poorly estimated. With greater redundancy in the population, it is easier to notice an individual poor tuning parameter estimate on some later day, allowing subsequent correction of the estimate and forestalling compounding errors. Sudden drift has an advantage not only because it introduces smaller errors under noisy observations from one day to the next (Fig. 3.e), but also because it can more effectively use of redundancy to fix error compounding (note the flattening gradient in Fig. 3.f and the better scaling with population size in Fig. 3.g), as from the perspective of the adaptive decoder a neuron with a low likelihood of its estimated tuning parameter is no different from a neuron that has spontaneously retuned.

Unsurprisingly, both gradual and sudden drift become easier to correct for under lower rates of drift, as the distribution of changes in tuning is narrower. Nevertheless, for the same population size and the same rate of drift, sudden drift is always easier to adapt to (Fig. 3.g). The learning that takes place within real neural populations may necessitate gradual changes, but supporting those gradual changes would require either larger populations or slower changes to achieve error rates equivalent to what is achievable with sudden tuning changes.

As a matter of rigour, we highlight that the definition for equivalence between sudden and gradual drift rates, which considers a long timescale over which the representation becomes completely scrambled, may be present an unfair disadvantage to gradual drift, whose Brownian motion-like exploration of parameter space can return to previous values. Using gradual drift to scramble the population beyond recognition in the same number of days as the sudden drift induces more error in a fixed decoder after the first day of drift (as visible in Fig. 3.e). However, even if we redefine the equivalence between sudden and gradual drift to cause identical degradation in a fixed decoder after the first day of drift (resulting in sudden drift models completely scrambling populations faster than gradual ones), we still find the same advantages of sudden of gradual drift (Fig. S2.c, S2.d).

### Is in vivo drift sudden or gradual?

Our computational results imply that neural populations with downstream readouts can facilitate drift compensation by making representational changes stark, especially in the case of fast drift rates. This would allow for populations to adjust faster or employ fewer neurons. Given the potential advantages, we expect that the statistics of in vivo drift reflect a preference for heavy-tailed drift.

To explore whether in-drift manifests drift suddenly or gradually, we return to our previous comparison of the distribution of pairwise correlations between neurons (as in Fig. 3.d). This has the advantage of being agnostic towards which features of environments, stimuli, or tasks a given population encodes, and therefore allows us to draw crude comparisons both between different in vivo datasets and between in vivo datasets and in silico models.

We reiterate that because both gradual and sudden drift will eventually scramble a population entirely, it becomes impossible to distinguish gradual from sudden drift over timescales that are long relative to the drift rate. In gradual models of drift, the correlation between a pair of neurons gradually increases or decreases as the neurons drift together or apart, while in sudden drift their correlation appears or disappears spontaneously. In the transient period in which it is possible to distinguish gradual from sudden drift, we model the distribution of these pairwise tuning changes as a mixture of two component Gaussian distributions, with one component describing noise-driven fluctuations and the other describing drift-driven changes in tuning (Fig. 4.a, Fig. 4.b). While simplistic, this mixture model allows us to qualitatively distinguish gradual from sudden drift. Under gradual drift, the component of this distribution ascribed to changes in tuning gradually separates from the component ascribed to noise. We can therefore identify gradual drift by inspecting the increase over time in the mean of component of the mixture model that corresponds to nonstationary changes (*µ*_2_), the behaviour of which can be approximated by an exponential decay towards a final value (Fig. 4.e). By contrast, under sudden drift, the two components are distinct from the outset, and there is therefore minimal change in the value of *µ*_2_ (Fig. 4.f).

**Figure 4.**
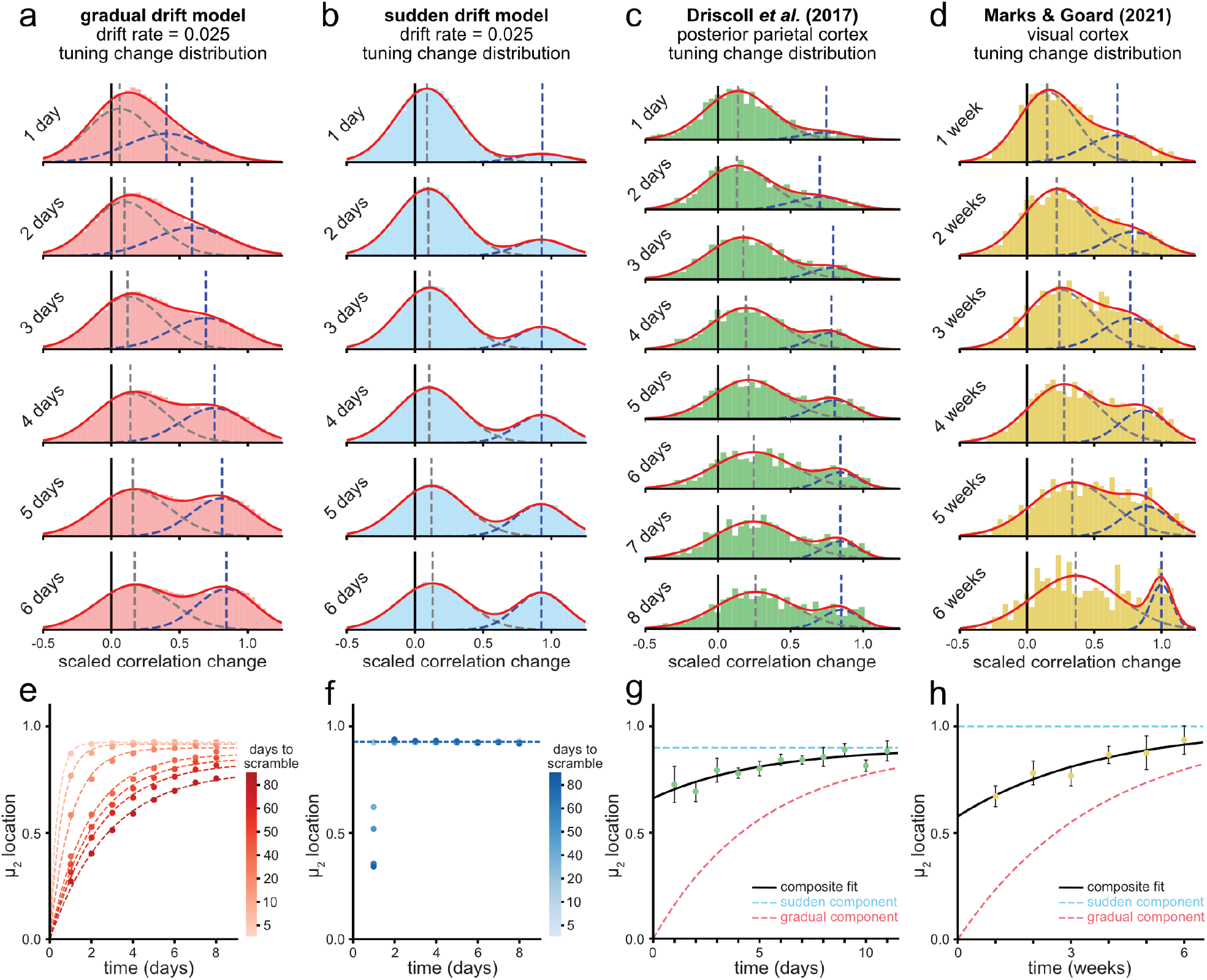
**(a)** Histograms showing the evolution of the distribution of changes in scaled correlations between neuron pairs in the gradual model of drift over the course of several days. The fit is a 2-component mixture of Gaussians, with each underlying component and its mean shown as dashed lines. **(b)** As in (a), but shown for the sudden model of drift at an equivalent drift rate and simple final-value fit. **(c)** As in (a), but shown for in vivo data from the Driscoll *et al*. (2017) dataset [7]. This comprises recordings from the posterior parietal cortex during performance of a T-Maze task. **(d)** As in (a), but shown for in vivo data from the Marks & Goard (2021) dataset [6]. This comprises recordings from the visual cortex during exposure to movies of naturalistic scenes. Location of the upper mean (*µ*_2_) in the mixture of Gaussians fit for the gradual drift model shown against time for a range of drift rates. The dashed line indicates a fit to the exponential approach approximation. **(f)** As in (e), but shown for the sudden model of drift and using a constant value approximation. **(g)** Location of the upper mean (*µ*_2_) in the mixture of Gaussians fit for the Driscoll *et al*. dataset. Error bars indicate +/-SEM as estimated via bootstrap. The fit is a weighted combination of gradual and sudden drift models, with each component rendered as a dashed line. **(h)** As in (g), but shown for the Marks & Goard (2021) dataset.

We first consider the dataset of recordings from the PPC in Driscoll *et al*. (2017) [7], where drift is observable on the timescale of days. On visual inspection, changes in the distribution of tuning do not fit neatly into the mould of either sudden or gradual drift (Fig. 4.f). While many changes appear to be mediated by large jumps in tuning, a gentle trend in the growth of *µ*_2_ implies that some component of the drift is also mediated by more gradual changes. We fit a simple weighted linear combination of our approximations for the sudden and gradual drift models for the evolution of *µ*_2_, and attribute relatively more of the changes in the distribution to sudden changes (Fig. 4.g).

Producing the distribution of correlation changes requires neuron pairs to remain intact between imaging sessions and to be have well-correlated activity on at least one of the sessions. The distribution in Fig. 4.f is an aggregate of all animals in the dataset. While some animals have more imaged neurons and more recording sessions (Fig. S3.a) and drift rates are not necessarily homogenous across animals, the proportion of representation of each animal within the data remains largely unchanged as a function of separation between sessions (Fig. S3.b), and therefore a change the ratios of which animals are represented is not responsible for changes in the shape of the distribution.

Next, we consider the dataset of recordings from V1 in Marks & Goard (2021) [6], where drift is much slower and observable on the timescale of weeks (Fig. 4.d). This too contains a mixture of sudden and gradual drift, but by comparison to the PPC data appears to contain a greater proportion of gradual drift (Fig. 4.h). As with the PPC data, this distribution is an aggregation over multiple animals with varying amounts of neuron pairs (Fig. S3.d and Fig. S3.d). The two datasets, which differ in rates of drift, differ also in the amount of drift which is mediated by gradual changes. For the PPC, the parameter balancing the ratio of sudden to gradual population changes in the composite fit attributes 73.6% (SEM=7.85%, estimated via bootstrap sampling of the tuning change distributions) of the changes in the population to sudden tuning changes, compared to 57.8% (SEM=8.35%) for the V1 fit. The population with the slower drift rate makes a greater proportion of tuning changes gradually (Welch t-statistic: 1.378, *p* = 0.084; pooled variance t-statistic: 1.829, *p* = 0.034).

## Discussion

In this work, we presented an argument and computational simulations in support of the advantages of populations mediating changes in representation through stark and sudden changes in the tuning of individual neurons. We are quick to highlight that the advantages of sudden changes in neuronal tuning do not preclude the existence of gradual changes: on the contrary, we demonstrate that it is possible to adapt to such gradual changes, and that at least some component of experimentally observed drift can be attributed to these gradual processes. However, we stress that to support such gradual changes, the population undergoing such changes must make concessions: it must either use a larger number of neurons and thus a higher degree of redundancy for its representations, or it must drift (or learn) more slowly, or it must rely on some external feedback system to compensate for drift.

Emerging ideas present drift, or at least some component of drift, as a direct consequence of the repeated application of learning rules implemented by neural populations [20, 21]. This viewpoint explains drift as a learning-driven walk exploring a degenerate parameter-space [22, 23]. Computational models of drift also assume gradual tuning changes at the level of individual neurons. This is either explicit, as in the modelling of drift as an exogenous phenomenon described by an Ornstein-Uhlenbeck process [18], or implicit, when using gradient descent-based learning rules to drive drift in a simulated population [23, 24] which result in similar Brownian motion-like drift in the parameter-space of populations [25, 26].

However, there are many statistical models of drift that can generate gradual drift in representation-space, and we argue that not all of these are functionally equivalent in terms of facilitating stable downstream readouts. Crucially, gradual drift in the tuning parameters of individual neurons may be disadvantageous. Instead, when distributions of tuning parameter changes are ‘heavy-tailed’, favouring sudden jumps in tuning rather than gradual shifts, it becomes easier, algorithmically, to distinguish nonstationary changes in representation from noisy observations.

We described a simple model process for heavy-tailed drift, a sudden retuning to a completely new parameter, to illustrate its advantages for distinguishing statistical fluctuation from change in tuning. In practice, other processes can perform this role better through distributions with even heavier tails. For example, upon retuning a neuron might be forced to adopt a new tuning parameter that is sufficiently different from its previous value. This ‘active avoidance’ of previous tuning values makes it even easier to distinguish statistical fluctuation from nonstationary change.

We acknowledge that there are myriad explanations for a discrepancy in sudden versus gradual tuning adjustments between the PPC and V1. In addition to the potential for physiological differences, these are different experiments recorded under different conditions during the performance of different tasks. Nevertheless, this preliminary consideration of in vivo data presents rudimentary agreement with the premise that a neural population can facilitate adaptation within downstream readouts by mediating representational changes through sudden changes in tuning, and that this advantage is more pronounced in situations involving higher drift rates. We find this corroborated by recent evidence of drift-like changes in tuning in CA1 hippocampus, in which representations make abrupt jumps in tuning on timescales visible within single recording sessions [27].

The relevance of the advantages of heavy-tailed changes in tuning extends beyond representational drift. Other nonstationary changes in the encoding of information of a population, such as the incorporation of new information during learning, face the same challenge of making themselves visible through the noise of day-to-day fluctuations. If we entertain the view that some component of drift is in fact caused by the repeated application to the same data of the same continual learning rules that incorporate new information, we open up the possibility of deriving constraints on the learning rules implemented by the brain through measurements of drift. While it is experimentally challenging to attribute new behaviours or newly learned information to changes in in vivo physiology, we speculate that–if measured–such changes would also demonstrate stark, heavy-tailed statistics.

Equally, we see future relevance of this work to in silico learning algorithms. Artificial neural networks are prone to drift when trained using gradient descent on a feed of continuously supplied data. This drift can, in turn, cause catastrophic forgetting for samples that have remained unseen for long periods of time. While techniques exist either to restrain this drift by explicitly incorporating penalty terms in loss functions [28] or by mitigating its consequences by actively identifying and utilising subspaces of parameter-space orthogonal to previously learned items [29], we are intrigued by the possibility that a combination of network redundancy and regularisation that discourages relatively small parameter changes might provide an implicit solution. By altering the dynamics of gradient descent to explore parameter-space by focusing, rather than distributing, updates to weights in a network, exposure to new samples disturbs previously learned representations in a way that discourages a tug-of-war between samples and instead encourages implicit identification of orthogonal subspaces of parameter-space.

The two key components of the implementation of our adaptive decoder are an accurate drift prior and a description of how neurons tune to a particular kind of stimulus in a population. While we do not claim that in vivo downstream populations explicitly perform Bayesian optimisation to account for changes in tuning, we do posit that it is physiologically plausible for a downstream population to make use of these key prior distributions in implementing an adaptive decoding strategy: the same brain regions across individuals of a species encode similar information using similar strategies and experience drift with similar statistics. Readouts across populations that connect regions therefore have access to reasonable priors about their upstream neural activity, which may be sufficient to implement adaptive decoding strategies.

We conclude with a call for greater scrutiny of the statistics of drift across brain regions. Drift rates measured in vivo vary considerably. Drift is observable in mere hours to days in highly plastic regions such as the hippocampus [3], but only on much longer timescales (weeks to months) in less plastic regions such as the visual cortex [6]. This relationship suggests a strong link between learning rates and drift rates, and additional factors such as the degree of novelty [9], the amount of experience [3], or potentially even the complexity of the task or richness of environment can modulate drift rate [21].

While drift is necessarily a gradual process at the scale of populations, this drift still permits a variety of underlying processes operating at the level of individual neurons. We predict that for the neural populations with the highest rates of drift and the highest rates of plasticity, changes in representation will take place through neurons performing sudden jumps away from previous tuning values in order to avoid ambiguity in downstream readouts.

## Methods

### Adaptive Decoding and Drift Simulations

#### Solving the optimisation for the gradual drift model

We use gradient ascent to solve for the most likely values of 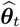.

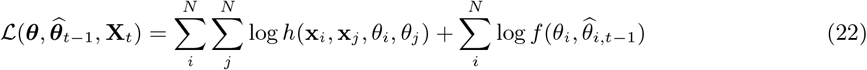

The left summation depends only on the neuron behaviour model and the right summation depends only on the drift model. In this simple example, we can find analytical forms of the Jacobian and the Hessian, which makes solving via gradient ascent faster. This makes use of two important properties of *h*. First, it is symmetrical over *i,j*:

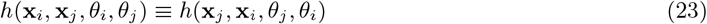

Second, in the special case when *i* = *j*, all choices of *θ*_*i*_ are equivalent.

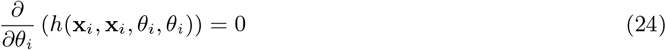

Combining these properties and taking the derivative of the log-likelihood with respect to a single parameter:

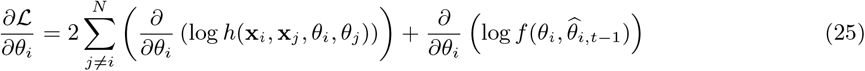

And then, to produce the Hessian:

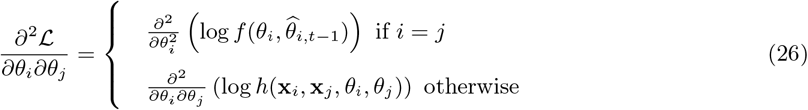

We initialise of the parameters for the gradient ascent is to use their values from the previous day, 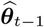. A complete expression and its derivation is provided in the supplementary material.

#### Exact solution to the optimisation for the sudden drift model

The discontinuity in *f*_sudden_ means that it is ill-conditioned for a gradient ascent-based solution applied to the entire parameter space. Instead, we consider a solution that decomposes the problem into smaller components that can be individually solved by gradient ascent (and, in the subsequent section, a technique for approximating this solution for large populations).

For each neuron, there are two possible cases: either the neuron keeps its tuning parameter, or it retunes. In the first case, trivially 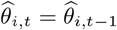. In the second case, the neuron needs a tuning parameter update *and*,.. should be withheld from use in estimating the parameters of other neurons this day, as *θ*_*i,t*−1_ is no longer accurate. To solve, we perform two steps:

- Infer which neurons have kept their tuning parameter intact
- Use this subset of neurons to adapt the parameters of the remaining neurons

Let the set *A* contain the neurons which have retained their tuning parameters. A neuron *i* is a member of this set if and only if *u*_*i*_ = 1, where:

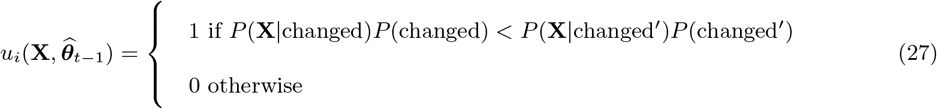

In the concrete example for the sudden drift model, the above condition can be written as:

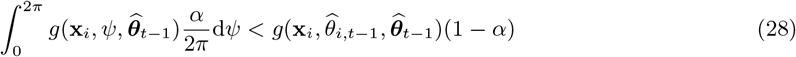

Where 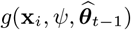 is the probability of the observations **x**_*i*_ given a tuning angle *ψ*.

This allows us to express the update rule as:

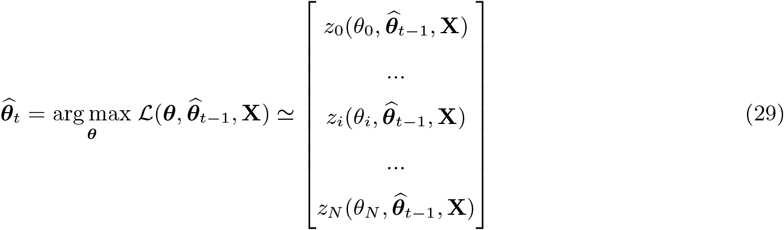

Where:

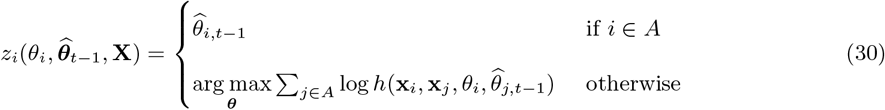

Noting that the use of the approximately equal symbol, as this solution discards the small amount of information available in the pairwise mutual tuning between neurons not in *A*.

#### Approximate solution to the optimisation for the sudden drift model

In practice, evaluating the condition for *u*_*i*_ becomes computationally intractible for large *M* due to resolution limitations of floating point number representations: it requires the product of very many small numbers. Circumventing this limitation in the typical fashion–a sum of logarithms–requires a closed form of the integral of *g*, which is not necessarily available. We instead choose to make an approximation in determining set membership of *A*.

We evaluate log 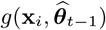 for each neuron in the population and are therefore able to rank each neuron in the population by conformity to expectation. For a drift rate parameterised by *α* and a number of neurons in the population *N*, the number of neurons that drifted is binomially distributed. We can select a probability threshold *q* (by using the CDF of the binomial distribution) that results in us excluding the bottom *Q* neurons from *A* by ranking.

The choice of *q* is a hyperparameter. If *q* is large (e.g. 0.95), many of the neurons excluded from *A* are likely to not require retuning. For small *N* and high drift rates (large *α*), this can exclude a sizeable portion of the population and reduce the accuracy of tuning parameter estimates. Conversely, if *q* is small (e.g. 0.5), neurons that require re-estimating will be included in *A* and not have their parameters updated. In all simulations shown in this work, we set *q* = 0.9.

This approximation has the potential to disadvantage an adaptive decoder compensating for sudden drift when compared to one compensating for gradual drift. Indeed, we observe that levels of noise high enough to make an adaptive decoder perform no better than a fixed decoder, this approximation leads to performance that is worse than rather than equivalent to a fixed decoder (as visible in Fig. 3.e and Fig. S2.c). However, given our choice of neuron population sizes *and* the continued outperformance of the sudden drift adaptive decoders, we take our use of this approximation to be immaterial to our conclusions.

#### Implementation of solvers

The gradient ascent components of these solutions were implemented using the `scipy`’s library’s `optimize.minimize` function (in the case of the multivariate optimisation for the gradual case) and the `optimize.minimize_scalar` function (in the case of the independent scalar optimisations for the sudden case) [30]. We note the additional following details:

- Solution bounds are set to −2*π* to 2*π*, and only remapped to the range −*π* to *π* after convergence
- The underlying solver method is based on `L-BFGS-B` [31]
- We use the analytical expression for the Jacobian as described in the supplementary material
- Exact parameters used for the solver, including increases to search depth and stricter tolerances from default usage of these solvers, are available in the code provided as supplementary material

### Tuning parameterisation-free drift characterisation

#### Distributions of correlation changes in vivo

For in vivo data, ‘correlation’ referes to the Pearson Correlation Coefficient. When visualising the distribution of correlation changes for experiments *T* sessions apart (in days for the simulated and Driscoll *et al*. (2017) dataset, in weeks for the Marks & Goard (2021) dataset), we include all session pairs separated by that interval. For example, if *T* = 2 in the Driscoll *et al*. (2017) dataset, we would include day 1 paired with day 3, day 2 paired with day 4, day 3 paired with day 5, and so on. Correlations between neuron pairs can only be evaluated within an individual animal. We aggregate all neuron pair correlations from all animals to produce the distributions. For both in vivo datasets, we select *C*_min_ = 0.5: this is high enough to cull spurious correlations (which mask the visibility of the effect of drift on the distribution), while low enough to include many pairs of neurons.

#### Mixture of Gaussian fits

We fit the distribution of correlation changes with a 2-component mixture of Gaussians. This is solved using the EM algorithm as implemented by the `scikit-learn` library [32]. For the in vivo data, we use bootstrapping (100 iterations per distribution) to fit many resampled distributions to produce the SEM used for the error-bars in Fig. 4.g and Fig. 4.h.

#### Modelling the movement of the upper mean of the mixture of Gaussians fit

In the case of sudden drift, we approximate fits to distribution of correlation changes as always being bimodal with a fixed upper mean (i.e. it is time independent).

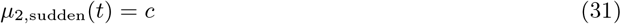

In the case of gradual drift, we approximate the upper mean of the mixture model fit to increase as a function of elapsed time, described by:

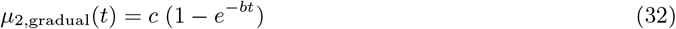

For the in vivo data, we fit a weighted mixture of the sudden and gradual models:

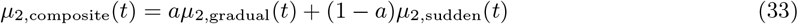

Noting the following constraints:

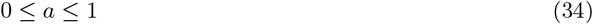

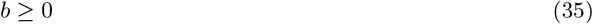

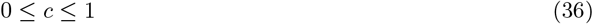

The parameters *a, b*, and *c* are found by minimising the mean squared error between this model of upper mean locations and its measured values.

### Drift in the posterior parietal cortex

#### Data Inclusion

The Driscoll *et al*. (2017) dataset comprises recordings from 5 animals (m01 through m05). Because our analysis relies on pairs of neurons being visible across sessions, the number of individual neurons strongly informs the quality of the data. Our analysis includes only neurons that are identified between sessions at the highest level of confidence as labelled in the dataset. Animals m02 and m05 have a distinctly lower total number of neurons identifiable between sessions (Fig. S3.a), experience a more rapid decline in the number of neurons identifiable between sessions as the number of days between those sessions increase, and exhibit a drop off over time in the number of identifiable neurons. Consequently, while we include all animals in our analysis, animals m01, m03, and m04 are more represented in the aggregated data. The proportion of which animals are representated in the data does not change substantially over time (Fig. S3.b).

#### Data Processing

To calculate the activity correlation between neuron pairs during each session, we first convolved spike times (from the original dataset as extracted by deconvolving Calcium traces) with a Gaussian window (standard deviation equivalent to 0.45 seconds). We then evaluated pairwise temporal correlation between these traces. We did not exclude neural activity during the interval of darkness between trials.

### Drift in the visual cortex

#### Data Inclusion

The Marks & Goard *et al*. (2021) dataset comprises recordings from 12 animals (Fig. S3.d). Mice 7 and 8 are omitted from our analysis as their imaging schedules deviate from the one used by the other animals. Mouse 2 is omitted from our analysis as, uniquely, its surgery provides two separate imaging windows. Our analysis includes only neurons labelled as having a registration quality of 3 or higher (unambiguously identified across sections). Our analysis makes use of only the experimental protocol which exposes animals to the movies of naturalistic scenes. As with the PPC dataset, the proportion of which animals are representated in the data does not change substantially over time (Fig. S3.e).

#### Data Processing

To calculate the activity correlation between neuron pairs during each session, we convolved the raw Calcium traces with a Gaussian window (standard deviation equivalent to 0.25 seconds). These traces were subsequently used as the basis for pairwise temporal correlation.

## Supplements

### Derivations

#### Expected correlation between neuron pairs

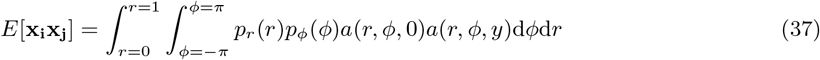

We note that *a*(*r, ϕ, θ*) is zero for |*ϕ* − *θ*| *> π/*2. Therefore, taking 0 ≤ *y* < *π*:

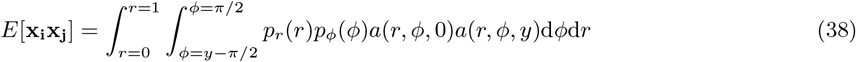

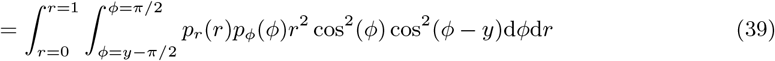

Noting that *r* and *ϕ* follow uniform distributions:

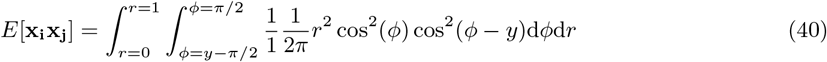

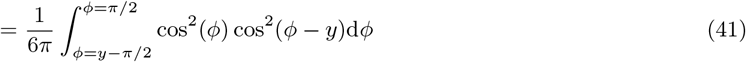

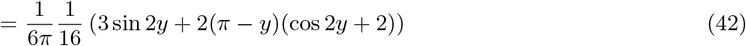

#### Jacobian for gradient ascent in gradual drift model

Then, the first partial derivative of log *h*:

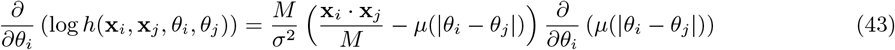

Where:

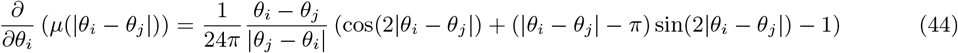

For the gradual drift prior, we have:

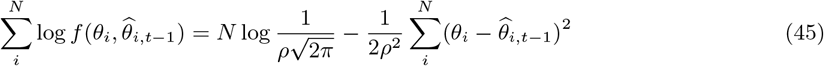

And the partial derivative:

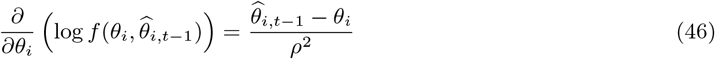

**Fig. S1:**
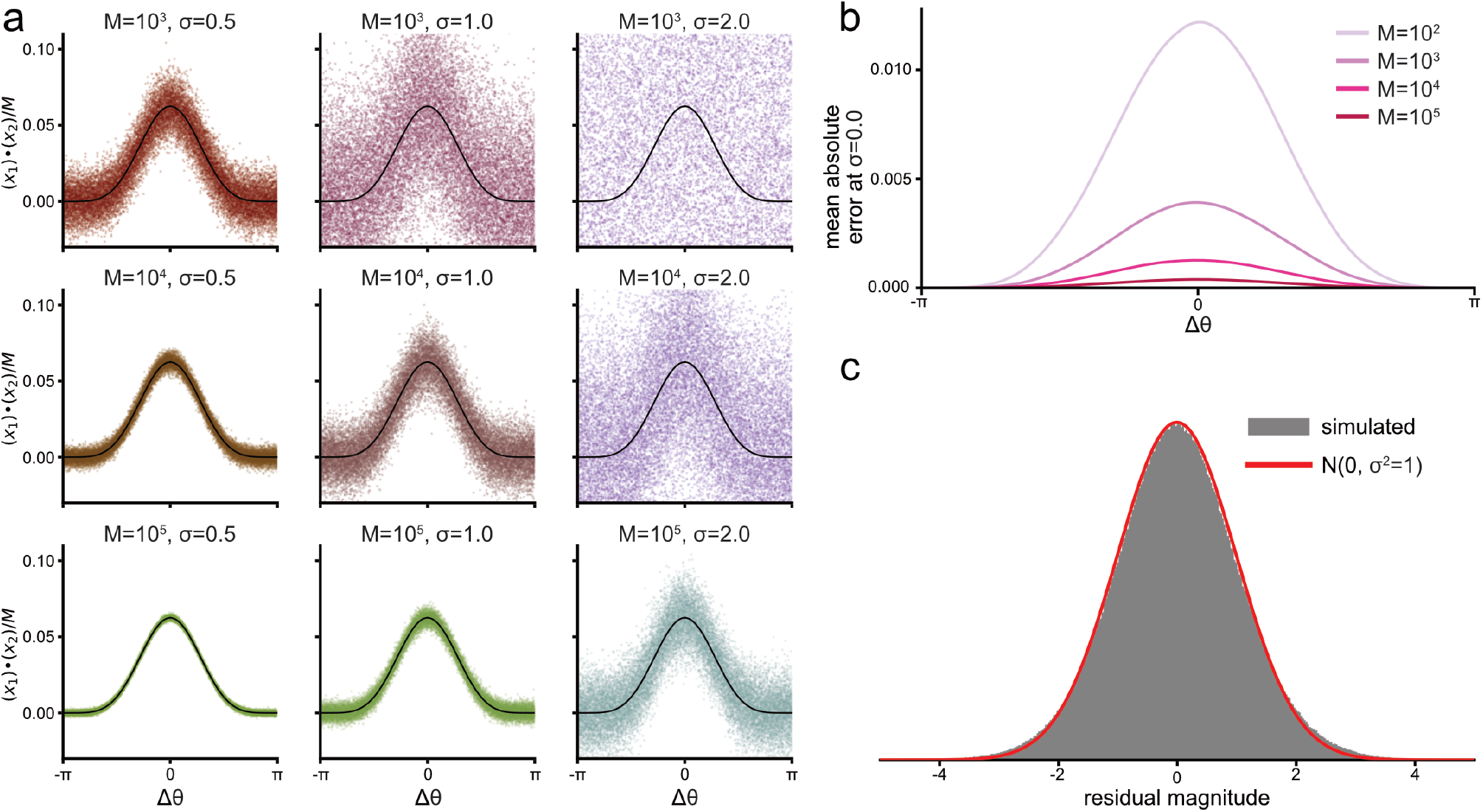
**(a)** Visualisations of the distribution of correlations between neuron pairs separated by an angular distance Δ*θ* for a variety of observation noise levels *σ* and total numbers of samples *M*. The black trace shows approximation for the expected value of the correlation. **(b)** The mean absolute error between the approximation for the expected value of the pairwise correlation and its actual value as computed by a Monte Carlo simulation, shown for in the absence of any observation noise (*σ* = 0) as a function of angular distance between the two neurons. **(c)** Distribution of the error residual, the difference between the expected value of the pairwise correlation and its actual value, in the presence of observation noise (*σ* = 1.0), presented with a comparison to the normal distribution, simulated at *M* = 15.

**Fig. S2:**
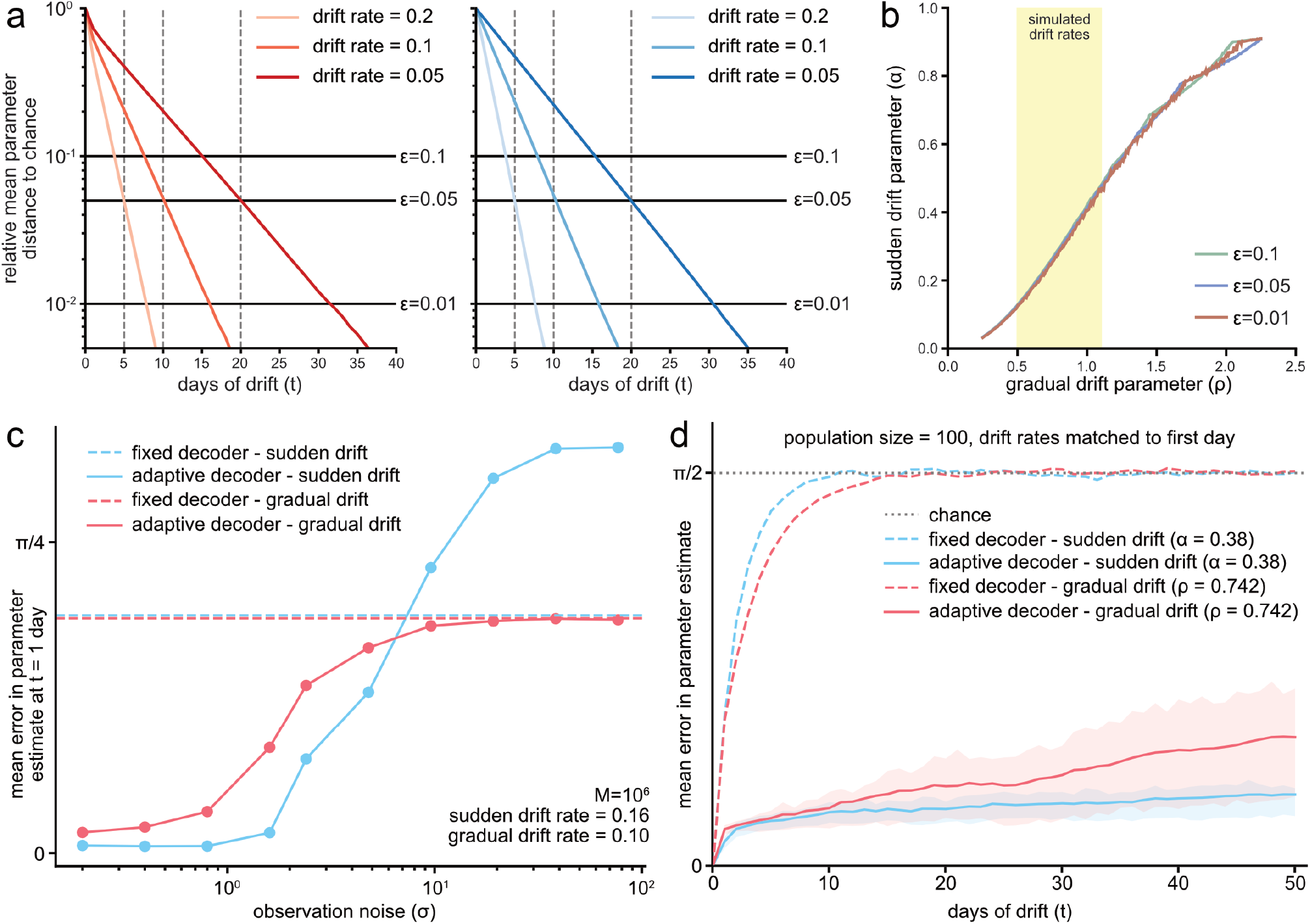
**(a)** The approach of tuning parameters to their chance values over the course of several days of drift, mean of 10^6^ simulations. Shown for the gradual drift model (left, red) and the sudden drift model (right, blue). Drift rates in the legend listed for *E* = 0.05. **(b)** The equivalence mapping between the gradual and sudden drift parameters for multiple values of *E*, highlighting the range of gradual drift rates used in the simulations of this article. **(c)** Error in tuning parameter estimates after the first day of drift as a function of the observation noise level *σ*, using alternative equivalence between gradual and sudden drift that induces the same amount of error in a fixed decoder after the first day of drift. **(d)** As in Fig. 3.f, but using the alternative drift equivalence.

**Fig. S3:**
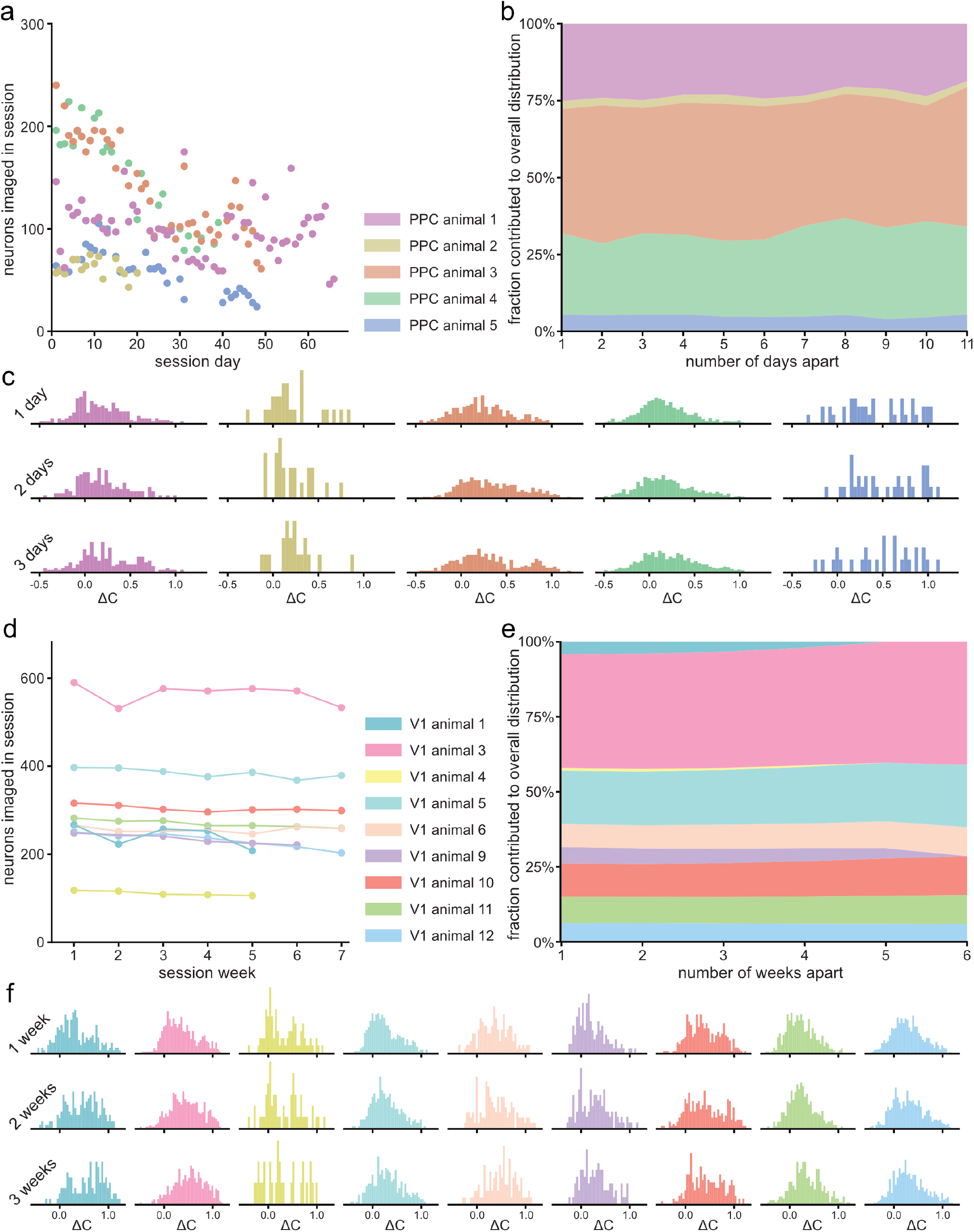
**(a)** The number of regions of interest (ROIs) labelled as neurons in each recording session for each animal of the Driscoll *et al*. PPC dataset. **(b)** The proportional contribution of each animal to the aggregated distribution of scaled changes in pairwise correlation. **(c)** Distributions of changes in pairwise correlation for the individual animals, shown for 1, 2, and 3 sessions apart. **(d)** The number of ROIs labelled as neurons in each recording session for each animal of the Marks & Goard V1 dataset. **(e)** As in (b), shown for the V1 dataset. **(f)** As in (c), shown for the V1 dataset.

